# Is there association between *APOE* e4 genotype and structural brain ageing phenotypes, and does that association increase in older age in UK Biobank? (N = 8,395)

**DOI:** 10.1101/230524

**Authors:** Donald M. Lyall, Simon R. Cox, Laura M. Lyall, Carlos Celis-Morales, Breda Cullen, Daniel F. Mackay, Joey Ward, Rona J. Strawbridge, Andrew M. McIntosh, Naveed Sattar, Daniel J. Smith, Jonathan Cavanagh, Ian J. Deary, Jill P. Pell

**Affiliations:** Institute of Health & Wellbeing, University of Glasgow, Scotland, UK; Centre for Cognitive Ageing and Cognitive Epidemiology, University of Edinburgh, Scotland, UK; Institute of Cardiovascular and Medical Sciences, University of Glasgow, Scotland UK; Department of Medicine Solna, Karolinska Institute, Stockholm, Sweden

**Keywords:** *APOE*, cognitive ability, ageing, brain MRI, grey matter, white matter

## Abstract

Apolipoprotein (*APOE*) e4 genotype is a purported risk factor for accelerated cognitive ageing and dementia, though its neurostructural substrates are unclear. The deleterious effects of this genotype on brain structure may increase in magnitude into older age. This study aimed to investigate in UK Biobank the association between *APOE* e4 allele presence vs. absence and brain imaging variables that have been associated with worse cognitive abilities; and whether this association varies by cross-sectional age. We used brain magnetic resonance imaging (MRI) and genetic data from a general-population cohort: the UK Biobank (N=8,395). We adjusted for the covariates of age in years, sex, Townsend social deprivation scores, smoking history and cardiometabolic diseases. There was a statistically significant association between *APOE* e4 genotype and increased (i.e. worse) white matter (WM) hyperintensity volumes (standardised beta = 0.088, 95 confidence intervals = 0.036 to 0.139, P = 0.001), a marker of poorer cerebrovascular health. There were no associations with left or right hippocampal, total grey matter (GM) or WM volumes, or WM tract integrity indexed by fractional anisotropy (FA) and mean diffusivity (MD). There were no statistically significant interactions with age. Future research in UK Biobank utilising intermediate phenotypes and longitudinal imaging hold significant promise for this area, particularly pertaining to *APOE* e4’s potential link with cerebrovascular contributions to cognitive ageing.

## Introduction

Variation at the *APOE* genetic locus is an established risk factor for Alzheimer’s disease (AD) (Lutz *et al.*, 2010), and cognitive decline in domains of memory, information processing speed and overall cognitive function (‘*g*’) (Lyall *et al.*, 2014). The e4 allele, which typically has a frequency of around 15% in Caucasian/European populations (Eisenberg *et al.*, 2010), is known as the ‘risk’ allele, vs. the neutral e3 allele (typical frequency 78%) and possibly protective e2 allele (frequency 6%). The *APOE* locus’s main function relates to lipid/cholesterol metabolism, which is pleiotropic for several biological functions including neuronal migration, axon guidance, and the clearance of amyloid beta plaques – which characterise AD - in the brain (Holtzman *et al.*, 2012).

There is evidence that the effects of *APOE* e4 variation on brain functioning increase across the lifespan, i.e. differences between e4 carriers vs. non-carriers in terms of cognitive ability become more pronounced with older age regardless of outright dementia (Schiepers *et al.*, 2011). Davies et al. (2015) reported data from the CHARGE consortium where in 53,949 European participants from 31 different cohorts (aged >45 years and non-demented), the strength of association between rs10119 (in the *APOE* genetic region) and a general factor of fluid cognitive function, increased linearly with the mean age of each cohort (Pearson’s *r* = −0.42, P = 0.022). Similarly, a report that included data from the Generation Scotland cohort (Marioni *et al.*, 2015) (n = 18,337) reported significant negative associations between *APOE* e4 allele presence and tests scores on Logical memory (standardized beta = −0.095, P = 0.003), and Digit symbol coding (standardized beta= −0.087, P = 0.004); however this was only significant in participants aged 60 years or over. We recently investigated potential *APOE* e4 genotype-by-age interaction on cognitive function in UK Biobank (n=~111k) (Lyall *et al.*, 2016); there were no statistically significant interactions; however there are concerns over the potential imprecision and reliability in the novel UK Biobank baseline cognitive tests, which means that result may be an underestimate of the potential true effect. Overall, recent research warrants further study into the potential structural brain substrates of the *APOE* e4 effect into older age.

It is of high importance to understand the neurobiological underpinnings of the potential age-related deleterious effects of *APOE* e4 on brain function. The integrity of connective WM tracts in the brain is associated with overall cognitive function (Penke *et al.*, 2012). The hippocampus is a known brain substrate of memory loss, one of the key symptoms of AD, and is the first site to show evidence of amyloid beta plaques (Reilly *et al.*, 2003). Total brain volumes, including total GM and WM volumes are significantly associated with changes in cognitive function (Brouwer *et al.*, 2014). Brain WM hyperintensities are a major substrate of age-related cognitive decline and cerebrovascular disease (Debette and Markus, 2010), and common in AD (Paternoster *et al.*, 2009).

Investigating associations between *APOE* genotype and various MRI markers is a topic of intense research, but studies involving structural neuroimaging parameters have not produced consistent results (Scarmeas and Stern, 2006). Whereas e4 carriers appear to be at increased risk of hippocampal atrophy (Manning *et al.*, 2014), worse WM microstructural integrity (Donix *et al.*, 2010; Fennema-Notestine *et al.*, 2011) and selective GM atrophy (Pievani *et al.*, 2009; Donix *et al.*, 2010; Fennema-Notestine *et al.*, 2011) in the context of pathological ageing (such as AD and mild cognitive impairment), there is less clarity with respect to MRI markers in non-pathological ageing. For example, null associations are reported for *APOE* genotype and cross-sectional hippocampal volume (Jack *et al.*, 1998; Killiany *et al.*, 2002; Schuff *et al.*, 2009; Ferencz *et al.*, 2013; Lyall *et al.*, 2013; Manning *et al.*, 2014), brain atrophy (Cherbuin *et al.*, 2008a), GM volume (Cherbuin *et al.*, 2008a) and some WM measures (Lyall *et al.*, 2014; Lyall *et al.*, 2015), though *APOE* e4 carriers exhibited greater WMH growth over a 3-year period in older age when compared to non-e4 carriers (Cox *et al.*, 2017). A meta-analysis (n = 8,917) reported that the *APOE* e4 genotype is associated with greater burden of MRI markers associated with cerebrovascular disease (cerebral microbleeds (Schilling *et al.*, 2013)). However, many prior studies possess low statistical power with which to detect subtle effects specifically in mid- and later-life, and thus it is unclear whether *APOE* status is unrelated to non-pathological brain ageing, or whether subtler differences exist, which prior studies have been unable to reliably quantify. Moreover, there is debate regarding the possible increasing importance of *APOE* e4 for brain structure with older age (Heise *et al.*, 2011).

The UK Biobank cohort is a large prospective cohort of whom around 12,000 currently have genetic and MRI data. This report aims to test for an effect of *APOE* e4 genotype carrier status on specific brain imaging phenotypes in UK Biobank, and whether that association interacts with age. The large sample N improves the potential reliability of association estimates herein, compared with smaller reports. We examined several specific brain imaging phenotypes based on a-priori hypotheses described above: left and right hippocampal volumes, total GM and WM volumes, WM hyperintensity volumes, and brain white matter tract integrity metrics.

It is hypothesized that for each brain phenotype, the association between *APOE* e4 genotype will be in the deleterious direction (i.e. lower scores for all except WM hyperintensities, for which higher scores are worse), and interact with age where the association becomes larger in older participants (on average). We assessed the phenotypes of: two latent measures of WM tract integrity derived from tract-specific FA (‘gFA’) and mean diffusivity (‘gMD’); total GM; total WM; total WM hyperintensity volume and hippocampal volumes (all in millimetres^3^). This will enable us to quantify, with considerable statistical power whether the effect of *APOE* e4 genotype on brain phenotypes of relevance to cognitive decline and AD is stronger in older age.

## Methods

### Study design and participants

The UK Biobank cohort is a large prospective cohort of 502,628 participants with phenotypic information. All participants attended one of 22 assessment centres from 2006 to 2010 where they completed a series of physical, sociodemographic, and medical assessments (Sudlow *et al.*, 2015). In 2014, MRI scanning of baseline participants began, and this is ongoing until around 100,000 participants have been scanned. As of November 2017, n=12,931 have MRI data derived by UK Biobank.

### Genetic data

UK Biobank genotyping was conducted by Affymetrix using a bespoke BiLEVE Axiom array for ~50,000 participants and the remaining ~450,000 on the Affymetrix UK Biobank Axiom array. All genetic data were quality controlled by UK Biobank (Bycroft *et al.*, 2017). The *APOE* e genotype is directly genotyped. Further information on the genotyping process is available (http://www.ukbiobank.ac.uk/scientists-3/genetic-data), including detailed technical documentation (https://biobank.ctsu.ox.ac.uk/crystal/docs/genotypingsampleworkflow.pdf). The two *APOE* e SNPs – rs7412 and rs429358 – were both in Hardy Weinberg equilibrium (P>0.05) assessed with PLINK V1.90 (Purcell *et al.*, 2007).

### MRI data

An average of four years after initial recruitment, a subset of UK Biobank participants also underwent head MRI on the same scanner at a single site (Cox *et al.*, 2016). The release of brain MRI data as of August 2017 is the subject of the current study. All brain imaging data used here was processed and quality checked by UK Biobank (Miller *et al.*, 2016; Alfaro-Almagro *et al.*, 2017) and available in the form of Imaging Derived Phenotypes (IDPs). Details on the UK Biobank imaging acquisition and processing including WM/GM and hippocampal segmentation, and on the WM diffusion processing, are freely available from three sources: the UK Biobank protocol: http://biobank.ctsu.ox.ac.uk/crystal/refer.cgi?id=2367 and documentation: http://biobank.ctsu.ox.ac.uk/crystal/refer.cgi?id=1977 and has also been described by Cox et al. (2016) and Miller et al. (2016). The total WM and GM variables were segmented automatically using FAST (Zhang *et al.*, 2001) and were normalised for skull size based on the T1 MR scan (as described in the open-access MRI protocol, pp.11; https://biobank.ctsu.ox.ac.uk/crystal/docs/brainmri.pdf). Total WM hyperintensity volumes were calculated based on T1 and T2 FLAIR, derived by UK Biobank using the Brain Intensity Abnormality Classification Algorithm (BIANCA) (Griffanti *et al.*, 2016). WM hyperintensity volumes were log-transformed here due to a positively skewed distribution.

FA and MD are commonly-derived WM tract integrity variables which describe the directional coherence and magnitude of water molecule diffusion, respectively. Water molecules tend to diffuse with greater directional coherence and lower magnitude when constrained by tightly-packed fibres (such as well-myelinated axons) as well as by cell membranes, microtubules and other structures; lower FA and higher MD are generally considered to reflect poorer / less ‘healthy’ white matter (Jones *et al.*, 2013). We constructed general factors of FA (gFA) and MD (gMD) using principal components analysis based on 22 tracts as described by Cox et al. (2016). These general measures reflect the high degree of shared microstructural properties across major white matter tracts in the brain; as found in this cohort, and various other groups (Penke *et al.*, 2012; Alloza *et al.*, 2016; Cox *et al.*, 2016; Telford *et al.*, 2017). Inspection of eigenvalues showed clear single unrotated factor solutions for gFA (eigenvalue = 12.83, r^2^ = 58%) and gMD (eigenvalue = 13.76, r2 = 63%). All brain MRI metrics were transformed into Z-scores based on the final analysis sample to ease interpretation.

### Covariates

UK Biobank derived a Townsend deprivation score for all participants immediately prior to baseline; calculated from data on car ownership, household overcrowding, owner-occupation and unemployment aggregated for postcodes of residence (Townsend, 1998). Higher Townsend scores equate to higher levels of area-based socioeconomic deprivation. Participants were asked during the baseline and MRI assessments about any previous or current cardiometabolic conditions that had been diagnosed by their doctor. Specifically, participants were asked whether their doctor had diagnosed each of myocardial infarction, angina, hypertension, diabetes or stroke. We defined CHD as either myocardial infarction (MI) or angina. Smoking was self-report as never, previous, or current smoker based on self-report; we simplified this into a binary ever (previous plus current) vs. never. We excluded participants that stated only ‘prefer not to answer' for disease and smoking variables: less than 1% of the sample.

### Exclusions

There were 12,662 participants with *APOE* e genotype and brain MRI data. We excluded participants with non-white British ancestry, self-report vs. genetic sex mismatch, putative sex chromosomal aneuploidy, excess heterozygosity, and missingness rate >0.1. This left n=11,065. We removed participants who reported a neurological condition at baseline or scan visit (~5% Supplementary Table 1); the inclusion of which could drive type-1 errors due to skewed results. We accounted for relatedness between participants by selecting one random participant for analysis from sets where two or more individuals were 1st cousins or closer. This left 8,395 participants for whom genotype frequencies of *APOE* e were e2/e2 n=48 (1%), e2/e3 n=1,032 (12%), e2/e4=208 (2%), e3/e3=4,960 (59.0%), e3/e4 n=1,958 (23%) and e4/e4 n=189 (2%). As a check, participants were split into age groups (under 50; 50 to 59; 60 and over): a chi square test showed no significant difference in e4 frequency (p=0.148).

### Analysis

We used an e4+ dominant model of e3/e4 and e4/e4 collated vs. e2/e2, e2/e3 and e3/e3 collated; e2/e4 is usually removed because it has potentially risk and protective alleles (Wisdom *et al.*, 2011). We elected for an e4 dominant (i.e. present vs. absent) rather than dose model (i.e. 0/1/2) because there were relatively few e4 homozygotes. We ran two linear regression models to examine associations between e4 genotype and each of the brain imaging parameters: gFA scores, gMD scores, left/right hippocampus, total GM, total WM and WM hyperintensity volumes in mm^3^, each transformed to Z-scores. For the ‘partially adjusted model’ we adjusted for age, genetic array, 8 principal components (for stratification) and sex. For the ‘fully adjusted model’ we then additionally adjusted for Townsend deprivation scores, ever vs. never smoking cigarettes, and self-reported doctor diagnosis of diabetes, hypertension and CHD. For interaction models we added the requisite e4*age interaction term. Finally, we re-ran analyses with added age^2^ and an e4*age^2^ interaction term to capture potentially curvilinear relationships.

We determined an alpha level of P <0.05 as nominal significance, and corrected for multiple comparisons with the false discovery rate (FDR) using a specialized program (Pike, 2011). We calculated statistical power using G*power 3 (Faul *et al.*, 2007). We conducted two additional post-hoc sensitivity analyses: firstly, we adjusted for X/Y/Z-coordinates of brain position during scanning, because it was available in only n=6,647 participants. Secondly, we adjusted the final results for baseline assessment centre (one of 22 across the United Kingdom), to reduce the possibility of any systematic procedural differences e.g. handling of blood samples (although note that the MRI scanning data we report on here was single-site).

A significant effect of e4 allele presence vs. absence would indicate mean level differences in the brain parameter of interest as a function of *APOE* e genotype (vs. absence) cross-sectionally. A significant interaction with age (or age^2^) would be consistent with a hypothesis of greater (i.e. accelerating) decline with increasing age, according to *APOE* e4 allele presence vs. absence.

## Results

Descriptive statistics are shown in Table 1, including chi square (for ordinal/categorical data) and ANOVA (for continuous data) tests of differences between *APOE* e4 presence vs. absence groups. There were 8,395 participants (mean age = 61.55, SD = 7.03, range = 46-73), of whom 3,953 were male (46.1%). G*Power 3 showed we had 99% power to find a standardised effect size of Cohen’s *d* = 0.1 (where 0.2 is considered small) for significant associations or interactions, based on our e4 present vs. absent sample sizes.

**Table 1:** Descriptive statistics

| *Demographic/lifestyle variables* | e4 absent<br>(N=6,040;<br>74%) | e4 present<br>(N=2,147;<br>26%) | Effect<br>size | P-value |
| --- | --- | --- | --- | --- |
| Age in years, mean (SD) | 61.66 (7.03) | 61.19 (7.04) | 0.07 | 0.009 |
| Gender, male N (%) | 2,916 (48.28) | 941 (43.83) | 0.04 | <0.001 |
| Townsend score, mean (SD) | -2.035 (2.58) | -1.98 (2.58) | 0.02 | 0.367 |
| Smoking history, ever N (%) | 2,249 (40.32) | 848 (39.55) | 0.07 | 0.536 |
| *Brain MRI values* |  |  |  |  |
| Left hippocampal volume<br>(mm<sup>3</sup>) | 3821.69<br>(468.75) | 3814.99<br>(446.79) | 0.01 | 0.613 |
| Right hippocampal volume<br>(mm<sup>3</sup>) | 3933.61<br>(480.72) | 3924.82<br>(478.75) | 0.02 | 0.521 |
| Normalised total grey<br>matter (mm<sup>3</sup>) | 796,681.00<br>(47,417.56) | 800,931.70<br>(46,699.26) | 0.09 | 0.002 |
| Normalised total white<br>matter (mm<sup>3</sup>) | 711,833.90<br>(40,033.25) | 713,252.30<br>(41,790.69) | 0.04 | 0.220 |
| White matter hyperintensity<br>vol. (mm<sup>3</sup>) [before log-<br>transform] | 3517.85<br>(4480.04) | 3710.22<br>(4657.41) | 0.04 | 0.157 |
| *Disease prevalence rates* |  |  |  |  |
| Coronary heart disease N, % | 194 (3.22) | 59 (2.75) | 0.01 | 0.287 |
| Hypertension N, % | 1,387 (22.99) | 492 (22.95) | <0.01 | 0.971 |
| Diabetes N, % | 298 (4.97) | 85 (4.00) | 0.02 | 0.067 |

Effect size = Cohen's *d* for continuous data; Cramer's *V* for frequency data.

There were no associations between *APOE* e4 and gFA, gMD (Table 2), left/right hippocampal volumes, total GM or total WM both normalised for skull size: indicating no mean-level difference in these brain parameters across the sample. There was a significant association between *APOE* e4 possession and greater WM hyperintensity volume (fully-adjusted standardised beta = 0.088, 95% CI = 0.036 to 0.139, P = 0.001). There were no significant e4*cross-sectional age interactions at P<0.05. Results were unchanged when we added an age^2^ term (i.e. non-linear), adjusted for X/Y/Z-coordinates of brain position during MRI scanning, or baseline assessment centre.

**Table 2:** Associations between *APOE* e4 presence and brain MRI variables (uncorrected for type-1 error).

|  | Partially adjusted |  |  |  | Fully adjusted |  |  |  |
| --- | --- | --- | --- | --- | --- | --- | --- | --- |
|  | *b* | Lower 95%<br>*CI* | Upper 95%<br>*CI* | *p* | *b* | Lower 95%<br>*CI* | Upper 95%<br>*CI* | *p* |
| <u>Association with *APOE* e4 presence</u> |  |  |  |  |  |  |  |  |
| gFA | -0.034 | -0.092 | 0.024 | 0.255 | -0.034 | -0.092 | 0.025 | 0.259 |
| gMD | 0.027 | -0.030 | 0.085 | 0.352 | 0.026 | -0.031 | 0.084 | 0.369 |
| Left hippocampus | -0.005 | -0.058 | 0.049 | 0.861 | -0.003 | -0.057 | 0.051 | 0.911 |
| Right hippocampus | -0.005 | -0.058 | 0.048 | 0.858 | 0.002 | -0.052 | 0.055 | 0.954 |
| Total GM (normalised) | 0.012 | -0.031 | 0.055 | 0.585 | 0.010 | -0.033 | 0.053 | 0.645 |
| Total WM (normalised) | 0.029 | -0.025 | 0.082 | 0.294 | 0.019 | -0.035 | 0.072 | 0.496 |
| Total WM hyperintensities | **0.093** | **0.041** | **0.145** | **4x10<sup>-4</sup>** | **0.088** | **0.036** | **0.139** | **0.001** |
| <u>*APOE* e4*age interaction</u> |  |  |  |  |  |  |  |  |
| gFA | <0.001 | -0.008 | 0.008 | 0.985 | <0.001 | -0.008 | 0.009 | 0.924 |
| gMD | 0.002 | -0.006 | 0.010 | 0.661 | 0.001 | -0.007 | 0.009 | 0.768 |
| Left hippocampus | -0.004 | -0.012 | 0.003 | 0.252 | -0.004 | -0.012 | 0.004 | 0.291 |
| Right hippocampus | -0.004 | -0.012 | 0.004 | 0.308 | -0.004 | -0.011 | 0.004 | 0.342 |
| Total GM (normalised) | 0.001 | -0.005 | 0.007 | 0.842 | 0.001 | -0.005 | 0.007 | 0.638 |
| Total WM (normalised) | -0.004 | -0.011 | 0.004 | 0.340 | -0.004 | -0.012 | 0.003 | 0.258 |
| Total WM hyperintensities | -0.001 | -0.008 | 0.006 | 0.816 | -0.002 | -0.009 | 0.005 | 0.614 |

Partially adjusted: *APOE* e4 presence vs. absence plus 8 genetic principal components, assessment centre, genetic array, age in years and sex. Bold typeface denotes $p < 0.05$ . Fully adjusted: (also) Townsend deprivation scores, self-reported diagnosed diabetes; hypertension; coronary heart disease, ever vs. never smoking. Abbreviations: GM grey matter, WM white matter, FA fractional anisotropy, MD mean diffusivity, *b* standardised beta, *CI* confidence interval, *APOE* apolipoprotein e, MRI magnetic resonance imaging.

As additional exploratory analyses, we analysed the 22 individual tract-specific FA/MD values identified as most sensitive to age (Cox *et al.*, 2016). There was one specific significant association, such that e4 carriers tended to have worse FA of the tract forceps major (fully adjusted standardised beta = −0.078, 95% CI = −0.138 to −0.018, P = 0.011). However, there were no e4 associations with MD, nor any interactions between e4 and age/age^2^ on FA/MD (P value range otherwise = 0.108 to 0.994; data available upon request). We re-ran the nominally significant e4 present vs. absent association with WM hyperintensity volumes, as an e4/e4 vs. non-e4 model: this was statistically significant (fully-adjusted model standardised beta = 0.217, 95% CI = 0.066 to 0.368, P = 0.005), with the caveat that there were relatively few e4 homozygotes (n=189).

The e4 allele present vs. absent association with WM hyperintensity volumes remained significant when correct for type 1 error with FDR (fully adjusted q-value = 0.038) but the association with tract forceps major FA did not (fully adjusted q-value > 0.05).

## Discussion

This report examined the potential association between *APOE* e4 genotype and brain imaging phenotypes of relevance to cognitive ageing and dementia. We have previously reported on the baseline cognitive data from UK Biobank (n=111,739) and reported generally no *APOE* e4*age interaction (Lyall *et al.*, 2016), although the sensitivity of the tests used is unclear because they were novel, very brief, and suffered a degree of ceiling/floor effects (Lyall *et al.*, 2016). We expected the brain imaging phenotypes to be more sensitive to age-related differences conditioned on *APOE* genotype. In terms of addition to the literature, this study is the largest single-site study of *APOE* e4 genotype and brain imaging metrics which are associated with cognitive ageing, and the principal findings are that e4 carriers had significantly increased WM hyperintensity volumes indicative of worse cerebrovascular health, but was not associated with other brain imaging metrics, and the association did not significantly vary by age.

With respect to our hypotheses, there was an association between *APOE* e4 genotype and significantly increased WM hyperintensity volume, such that e4 carriers exhibited 0.09 SDs greater load than non-carriers, and e4/e4 homozygotes around 0.22 SDs; this reinforces findings in smaller meta-analysis (e.g. n=4,024) (Schilling *et al.*, 2013). One limitation of our data is that the exact mechanisms which lead to rarefication of white matter tissue, is unclear, and may be due to different causes in different people (Wardlaw *et al.*, 2013). The lack of a significant interaction with age provided no evidence that this effect was stronger at older ages; i.e. it was not the case that age and e4 genotype were synergistic. We found no significant associations or age interactions for *APOE* e4 with gFA, gMD, total GM, WM, or hippocampal volumes. The cross-sectional age range in the current sample, where most participants were aged 50 to 70 years, may limit our ability to find a significant interaction with age.

We found no significant interaction between cross-sectional older age, *APOE* e4 genotype and worse WM tract integrity (indexed by gFA/gMD) in around 8,000 middle to older-aged adults. This is an unusually large non-consortium, single-scanner imaging genetics study, which allowed relatively high statistical power to reliably detect small effects; however we did not find any significant *APOE* e4 genotype-by-age interactions.

During exploratory analyses, we did find one relatively novel *APOE* e4 association with a specific tract FA value – namely in the tract forceps major. It is interesting that this tract was not assessed in the previously largest *APOE* e4/MRI study, using the Lothian Birth Cohort 1936 (n=650) (Lyall *et al.*, 2014). This demonstrates the importance of assessing as many specific WM tracts as can be reliably identified in imaging studies of *APOE* e genotype, and may indicate some differential sensitivity of some tracts (Lyall *et al.*, 2014; Cox *et al.*, 2016). This association did attenuate when corrected for type-1 error with FDR, however, and may therefore reflect a spurious finding. It is however worth noting that brain imaging/diffusion tensor phenotypes are usually strongly correlated (Cox et al., 2016) and this can make adjustment for type-1 overly cautious in some circumstances; independent replication in different cohorts is therefore advised (Pike, 2011).

Recent cross-sectional studies investigating non-demented cognitive ability and *APOE* genotype have reported an interaction whereby older age magnifies the association estimate, e.g. Marioni et al. (2015) and Davies et al. (2015). Our study aimed to provide a potential neural substrate to this effect where declines in brain structure and integrity were greater in participants with *APOE* e4 genotype. We generally found no significant *APOE* e4 genotype-by-age interactions on total GM, total WM, left or right hippocampal volumes or total WM hyperintensity volume. This is in line with some other findings; Lyall et al. reported that of six relatively large (N range 198 to 949) studies which examined *APOE* e4 genotype and hippocampal volumes, only two were statistically significant at P<0.05 (Lyall *et al.*, 2013). Generally, there is no evidence of a main effect of *APOE* e4 on GM volume in non-demented adults (Cherbuin *et al.*, 2008b). In terms of WM hyperintensities, one meta-analysis reported no significant association (n = 7,351) however most studies were of observer-rated WH hyperintensities e.g. Fazekas scores (Paternoster *et al.*, 2009) while another did report an association with continuous WM hyperintensity volumes (n = 8,917) (Schilling *et al.*, 2013) (standardised beta = 0.047, 95% CI = 0.0006 to 0.094, P = 0.05). Our results are therefore mostly in line with a generally null association between *APOE* e4 and cross-sectional brain volumes in middle age (around 60 years), but not WM hyperintensity volumes and this may suggest that at least part of *APOE* e4’s contribution to worse cognitive ability is via a cerebrovascular-type pathway.

In terms of limitations, this study tested for a cross-sectional effect of increasing age on brain WM microstructure and structural morphology; a longitudinal within-participants design may be more informative because it would minimize cohort effects associated with being born in different time periods (Anstey *et al.*, 2003). UK Biobank will ultimately conduct longitudinal scanning in ~10,000 participants (Miller *et al.*, 2016). The participants in the current study were of generally good health; we excluded participants who self-reported neurodegenerative diseases or those likely to affect the brain. However these diagnoses are not validated medically and we therefore cannot be certain of their accuracy; a recent analysis of self-reported rheumatoid arthritis in UK Biobank showed that only around half of participants that reported the chronic illness were on some sort of medication, which puts the validity of the diagnoses into question (Siebert *et al.*, 2016).

There may be a degree of selection bias in the current sample, where primarily healthier older participants are likely to respond positively to invitation and attend assessment. It is also possible that this occurs across the whole sample with regards socioeconomic status, whereby more middle-class, professional people are liable to participate; we attempted to correct for this with the Townsend scores however these are proxies for deprivation rather than being based on individual-level data. Generally, the rate of participants with exclusionary diseases like stroke were similar to in the full UK Biobank cohort of approximately 500,000 – suggesting little bias in that regard between participants that attended baseline assessment from 2006-2010, vs. MRI scanning more recently. In terms of deprivation the mean Townsend score in this MRI sample was around −2.0 vs. −1.3 for the full baseline cohort; i.e. the current sample was slightly less deprived. There may also be a degree of survival bias where the *APOE* e4 by age interaction is underestimated because the older people with a more deleterious effect of e4 are more likely to be missing (Heffernan *et al.*, 2016): we saw no significant difference in e4 frequency by age group (<50; 50-59; ≥60) although there was a small but significant overall continuous effect where the e4 carriers were on average around half-a-year younger.

In terms of future research, UK Biobank will ultimately include biomarker and serum lipid data (e.g. high/low density lipoproteins). These data may be used to inform part of the causal chain from *APOE* genotype (a lipid transporter gene) to cognitive/brain phenotypes, including using Mendelian randomization methodologies (Bowden *et al.*, 2015). While this report includes MRI and genetic data on around 10,000 participants from UK Biobank, ultimately around 100,000 will have data on these by around 2023 (Miller *et al.*, 2016); this will permit even greater statistical power in future with which to detect such effects with greater reliability still.

## Ethical statement

This study was conducted under generic approval from the NHS National Research Ethics Service (approval letter dated 17th June 2011, ref 11/NW/0382). The authors have no competing interests to report.

## Acknowledgements

This research has been conducted using the UK Biobank resource; we are grateful to UK Biobank participants and research team members who collected the data, administered the data release and to the UK Biobank MRI team who processed and checked the MRI variables. This research was conducted using UK Biobank application 17689. Thanks to Prof. Cathryn Lewis for helpful suggestions during the UK Biobank Psychiatric Genetics conference call on 11.10.16. UK Biobank was established by the Wellcome Trust medical charity, Medical Research Council (MRC), Department of Health, Scottish Government and the Northwest Regional Development Agency. It has also had funding from the Welsh Assembly Government and the British Heart Foundation. Cox is supported by an MRC grant (MR/M013111/1) and the work was undertaken within the University of Edinburgh Centre for Cognitive Ageing and Cognitive Epidemiology (http://www.ccace.ed.ac.uk), part of the cross council Lifelong Health and Wellbeing Initiative (G0700704/84698 and MR/K026992/1). Funding from the Biotechnology and Biological Sciences Research Council (BBSRC), Engineering and Physical Sciences Research Council (EPSRC), Economic and Social Research Council (ESRC) and MRC is gratefully acknowledged.

## Authors’ contributions

All authors contributed substantively to this work. DML and SRC were involved in study conception and design. DML, SRC and JW were involved in data organisation and statistical analyses. JPP NS JC DJS IJD were involved in application to UK Biobank and data coordination. DML with assistance from SRC, RJS, BC, DJS, JC and LML drafted the report. All authors were involved in reviewing and editing of the manuscript and approved it.

---

**Supplementary Table 1:**
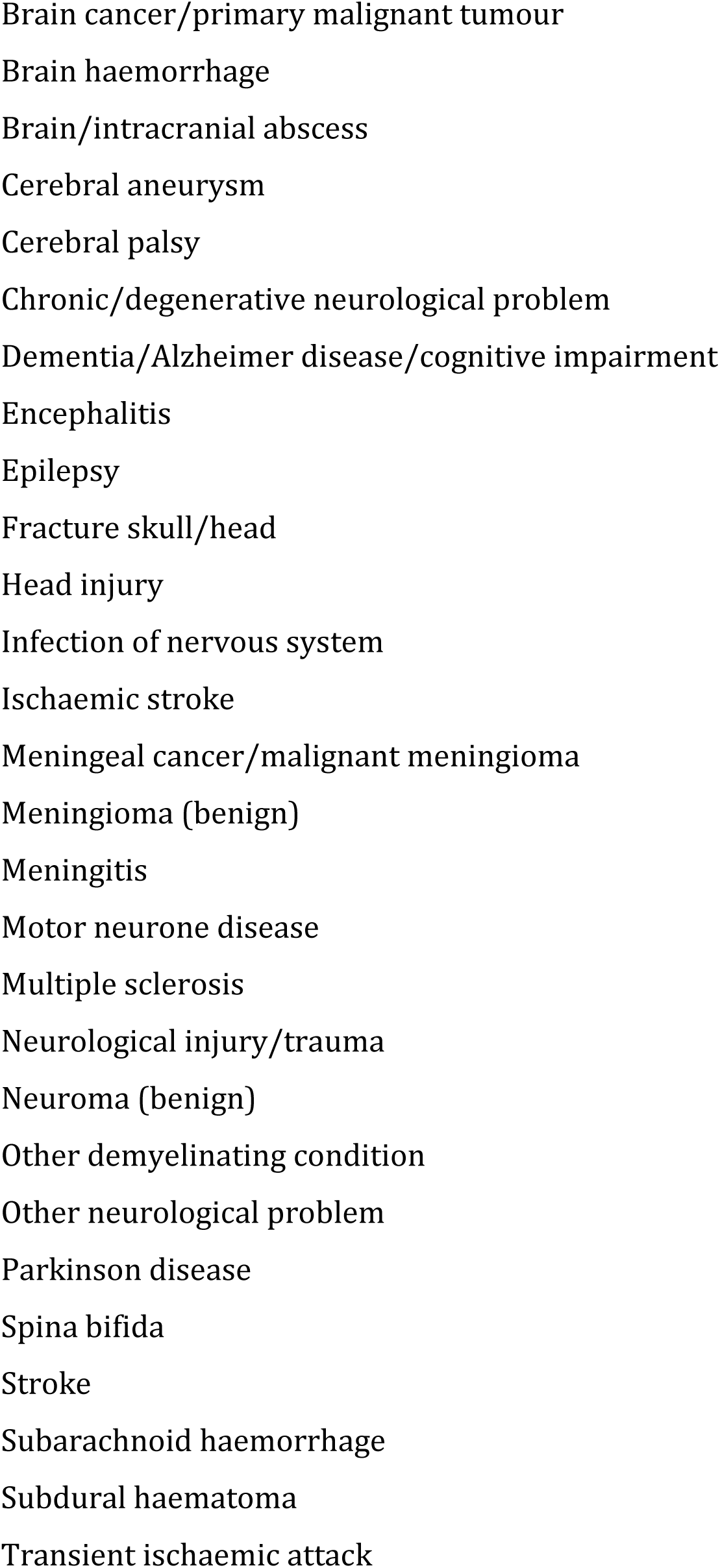
Excluded (self-reported) diseases

